# Lipid Desaturation Regulates the Balance between Self-renewal and Differentiation in Mouse Blastocyst-derived Stem Cells

**DOI:** 10.1101/2022.07.28.501839

**Authors:** Chanchal Thomas Mannully, Reut Bruck-Haimson, Anish Zacharia, Paul Orih, Alaa Shehadeh, Daniel Saidemberg, Natalya M Kogan, Sivan Alfandary, Raphael Serruya, Arie Dagan, Isabelle Petit, Arieh Moussaieff

## Abstract

Stem cells are defined by their ability to self-renew and to differentiate, both shown in multiple studies to be regulated by metabolic processes. To decipher metabolic signatures of self-renewal in blastocyst-derived stem cells, we compared early differentiating embryonic stem cells (ESCs) and their extra-embryonic counterparts - trophoblast (T)SCs to their self-renewing counterparts. A metabolomics analysis pointed to the desaturation of fatty acyl chains as a metabolic signature of differentiating blastocyst-derived SCs via the upregulation of delta-6 desaturase (D6D; FADS2) and delta-5 desaturase (D5D; FADS1), key enzymes in the biosynthesis of polyunsaturated fatty acids (PUFAs). The inhibition of D6D or D5D by specific inhibitors or SiRNA retained stemness in ESCs and TSCs, and attenuated endoplasmic reticulum (ER) stress-related apoptosis. D6D inhibition upregulated stearoyl-CoA desaturase-1 (Scd1) in ESCs, essential to maintain ER homeostasis. In TSCs, however, D6D inhibition downregulated Scd1. TSCs show higher *Scd1* mRNA expression and high levels of monounsaturated fatty acyl chain products in comparison to ESCs. Addition of oleic acid – the product of Scd1 (essential for ESCs), to culture medium, was detrimental to TSCs. Interestingly, TSCs express a high molecular mass variant of Scd1 protein, hardly expressed by ESCs. Taken together, our data point to lipid desaturation as a metabolic regulator of the balance between differentiation and self-renewal of ESCs and TSCs. They point to lipid polydesaturation as a driver of differentiation in both cell types. In contrast, mono unsaturated fatty acids (MUFAs), known to be essential for ESCs are detrimental to TSCs.

## INTRODUCTION

The regulation of cell viability by metabolic perturbations has been well-documented for a century. The set of small molecules (“metabolome”) has accordingly emerged as an important means for a detailed examination of cell state. While most studies describing the metabolic regulation of cell state and cell viability, have been carried out on cancer cell lines, recent studies strongly suggest that metabolic perturbations also play major roles in stem cell (SC) biology.

The balance between self-renewal and differentiation of SCs of the blastocyst is critical for their function *in vitro* and *in vivo*, and enables the concerted development of the embryo. The blastocyst comprises an inner cell mass (ICM; consisting of the pluripotent epiblast, which generates the future embryo, and the extra-embryonic primitive endoderm), and the trophectoderm (1). During morula compaction and the formation of the blastocyst, cells undergo the first differentiation into ICM and trophoblast (2). Cells that are isolated from the ICM can be cultivated as embryonic (E)SCs, defined by their ability to differentiate into all adult cell types (3), hence pluripotent (P)SCs. Cells isolated from the trophectoderm give rise to the placenta (4, 5), and can be cultivated as trophoblast (T)SCs. Importantly, the low accessibility of early embryos, the paucity of embryonic material, and the technical difficulties in direct experimental manipulation *in vivo*, underscore the necessity of cell models for the study of early development (6). This is especially true for metabolic analyses, as there is no amplification of signal in such analyses, thereby requiring a relatively large amount of cells per sample.

Research during the last decade by several groups, including ours, has uncovered metabolic signatures of SC potency (7–12), and roles played by metabolic shifts in the molecular circuitry that maintains stemness (7, 10, 11, 13–34). Metabolic perturbations also regulate self-renewal in pluripotent and adult SCs (13, 31, 35–39). Taken together, these studies suggest that the metabolic status of SCs is critical for the balance between self-renewal and differentiation in SCs.

We previously demonstrated a shift in glycolytic flux in PSCs upon their exit from pluripotency, driving differentiation via modification of histone acetylation (7). While shifts in central metabolism (7, 14, 19, 20, 24) and amino acid metabolism (11, 27) have been suggested to regulate the turnover and differentiation of SCs, information on the roles of lipid pathways in SC biology is still lacking. Nevertheless, lipids represent a major energy source during early development (40), and their content was reported to regulate the self-renewal of human ESCs (41). In particular, phospholipids are involved in PSC differentiation, as CDP-ethanolamine pathway synthesis of phosphatidylethanolamine is required at the early stage of reprogramming to PSCs (42).

In a seminal study that provided an unbiased view of ESC metabolism vs differentiated cells, Yanes and colleagues found highly unsaturated metabolite structures in undifferentiated ESCs (9). Importantly, however, inhibition of polyunsaturated fatty acid (PUFA) synthesis promoted pluripotency, suggesting that lipid polydesaturation may have a unique non-redox role in the differentiation of ESCs. Further support to the notion that the desaturation of fatty acyl chains plays critical roles in SC metabolism was given by Ben-David and colleagues, who demonstrated that stearoyl-CoA desaturase (Scd)1, also called Δ-9-desaturase, is essential for ESC survival (37). In mammalian cells, Scd1 is a rate limiting enzyme in the conversion of saturated fatty acids (SFAs) to monounsaturated fatty acids (MUFAs).

Here, we sought to reveal the metabolic events that take place upon early differentiation of blastocyst-derived SCs, and in particular, the ICM-derived ESCs and their extra-embryonic equivalents, the trophectoderm-derived TSCs. We found higher abundance of polyunsaturated acyl chains in both cell types upon early differentiation. The inhibition of Δ-6-desaturase (D6D, or FADS2) or Δ-5-desaturase (D5D, or FADS1) attenuated differentiation in both cell types, resulting in increased self-renewal. Interestingly, Scd1 inhibition showed opposite effects on cell viability in ESCs and TSCs, revealing distinct metabolic requirements between these two embryonic SC populations.

## RESULTS

Our previous work demonstrated a metabolic shift in ESCs upon the exclusion of growth factors that retain pluripotency, causing early differentiation (7). Following the same approach, we now sought to unveil common and distinct metabolic regulatory mechanisms of the balance between differentiation and self-renewal in ESCs and TSCs. For this aim, we allowed ESCs or TSCs to differentiate, and compared their metabolic profiles to the profiles of their self-renewing counterparts.

ESCs were cultivated in feeder free conditions for at least 3 passages. 24 hours after seeding, we initiated spontaneous differentiation, by exclusion of LIF and 2i from feeder free culture medium, and collected cells for a metabolomics analysis 48 hours after medium change. For undifferentiated control cells, ESCs of the same experimental lot were maintained in ESC culture medium that contained LIF and 2i and was replaced with fresh medium containing LIF and 2i. We then compared the early differentiating ESCs to their undifferentiated counterparts. The colonies of early differentiating ESCs showed a non-condensed morphology with unclear borderlines (**Figure 1A**), suggesting their exit from pluripotency. This observation was corroborated by decreased expression of the pluripotency markers *Nanog* and *Sox2* (**Figure 1B**). No significant change was noted in the expression of *Oct4*, known to be downregulated more slowly upon differentiation (43). Partial least squares regression discriminant analyses (PLS-DAs) of the LC-MS-derived metabolic profiles following quantile normalization showed discrete metabolome of the early differentiating ESCs (**Figure 1C**; R^2^=0.89, Q^2^=0.69 for blastocyst derived ESCs; R^2^=0.93, Q^2^=0.84 for TNGA *Nanog* ESCs).

**Figure 1.**
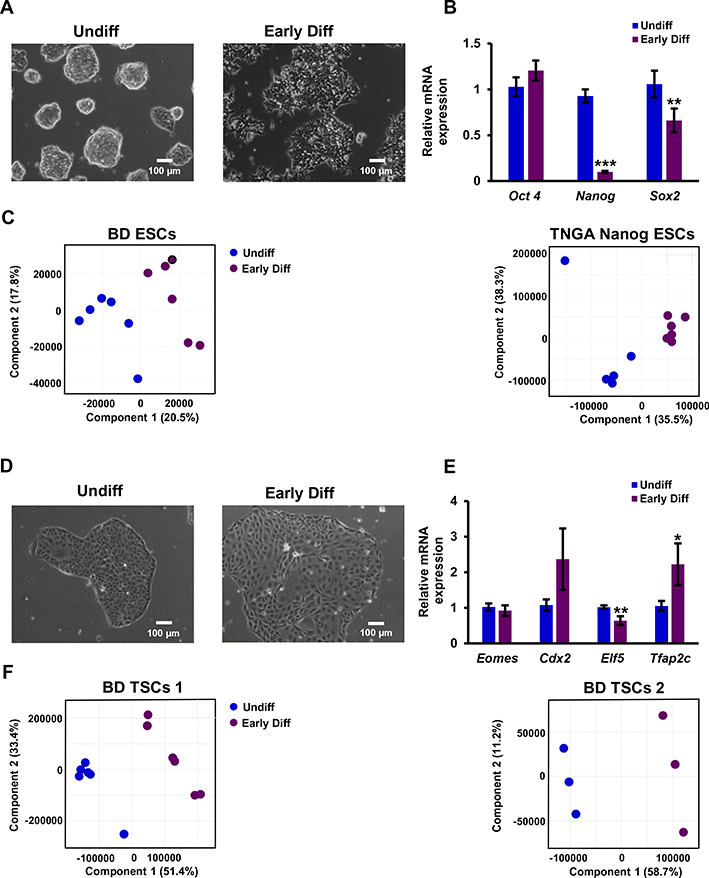
The metabolism of blastocyst-derived SCs shifts upon early differentiation. ESCs and TSCs (0.5 × 10^6^ cells per sample) were seeded in the culture media commonly used for their maintenance (Materials and Methods). After 24 hours, medium was changed to fresh medium with (“Undiff”) or without (“Early Diff”) factors that support self-renewal. ESCs were allowed to differentiate by the exclusion of LIF and 2i from cell medium. The colonies of early differentiating ESCs show a non-condensed, non-organized morphology (A). Early differentiating ESCs of the experimental lot used for the metabolomics analysis were evaluated for the expression of pluripotency markers by RT q-PCR (B). Following quantile normalization of the data, partial least squares discriminant analyses (PLS-DA) of early differentiating vs undifferentiated ESCs was performed. R^2^=0.89, Q^2^ = 0.69 for hetero agouti (HA) blastocyst derived ESCs (BD ESCs); R^2^ = 0.93, Q^2^ = 0.84 for TNGA *Nanog* ESCs (C). TSCs were allowed to differentiate by the exclusion of FGF4, TGFβ and heparin from cell medium. The colonies of early differentiating TSCs show a non-condensed non-organized morphology (D). The expression of TSC marker genes upon early differentiation was evaluated by RT q-PCR (E). Following quantile normalization of the data, PLS-DA of early differentiating vs undifferentiated TSCs. R^2^ = 0.99, Q^2^ = 0.98 for blastocyst derived (BD TSC 1, R^2^ = 0.99, Q^2^ = 0.91 for BD TSC 2 (F). Values are the mean ± SEM. n=6. *, P < 0.05; *, P < 0.01; **.

In a similar manner, we cultivated TSCs in feeder free medium (See Materials and methods) for 3 passages, and then excluded FGF4, TGFβ and heparin from cultivation medium, to initiate spontaneous differentiation. The medium of TSCs of the same experimental lot was replaced with fresh TSC culture medium that contained FGF4, TGFβ and heparin, and they were used as undifferentiated control. Early differentiation was confirmed by morphology (**Figure 1D**) and the expression of TSC marker genes. Specifically, the TSC marker gene *Elf5* was downregulated and the trophoblast lineage marker *Tfap2c* was upregulated (**Figure 1E**) (44). The expression of *Eomes* and *Cdx2*, two more TSC markers, did not significantly change. We compared the metabolome of TSCs after 48 hours of differentiation to that of their undifferentiated counterparts, and found a full separation: *Oct4*-GFP M2rtTA homo BL6 TSCs (blastocyst derived TSC1; “BD TSC 1”; R^2^=0.99, Q^2^=0.98), and *Oct4*-GFP M2rtTA hetero agouti (“BD TSC 2”; R^2^=0.99, Q^2^=0.91) (**Figure 1F**).

After demonstrating the metabolic shift in ESCs and TSCs in standard cultivation media, we wished to study these shifts in ESCs and TSCs maintained in similar defined cultivation medium. As TSCs have strict requirements for medium composition, and cannot be maintained in conditions suitable for ESCs, we maintained ESCs and TSCs in Tx basic medium, suitable for TSC cultivation (See Materials and Methods). We supplemented Tx medium with FGF4, TGFβ and heparin for TSCs, or with LIF and 2i for cultivation of ESCs. 24 hours following seeding, we allowed SCs to differentiate by excluding the above factors from cultivation medium for 48 hours. The medium of SCs of the same experimental lot was replaced with fresh ESC/TSC culture medium as above, and they were used as undifferentiated control. We then compared the metabolic composition of early differentiating SCs to that of undifferentiated cells. A PLS-DA demonstrated a distinct common metabolic shift along the second principal component (PC2), separating self-renewing SCs from early differentiating SCs (**Figure 2A**). To reveal the metabolites responsible for the separation, we performed a variable importance in projection (VIP) analysis. Of the 132 VIP>1 list of metabolic features, we noted a large number of phospholipids (39). Of these, MS/MS experiments confirmed the putative identifications of 19 phospholipids with more than two double bonds (polyunsaturated) in their acyl chains. Notably, phospholipids with polyunsaturated acyl chains showed higher abundances in early-differentiated TSCs and ESCs (**Figure 2B; Table S1**). The acyl chain composition of phospholipids is an accurate and sensitive measure of fatty acid incorporation into the cells (45). Accordingly, we assumed that the total content of polyunsaturated lipid species in the early differentiating SCs is increased. In agreement with this premise, we measured higher concentrations of total arachidonic acid in the early differentiating ESCs with no change in the levels of linoleic acid. (**Figure 2C-D**). We next wished to determine the enzymes responsible for the shift in lipid desaturation in the early differentiating SCs. Polydesaturation in animals requires sources for lipids with two double bonds, primarily linoleic and α-linolenic essential fatty acids. Delta-6-desaturase (D6D; fatty acid desaturase 2; FADS2) and delta-5-desaturase (D5D; FADS1) catalyze the synthesis of PUFAs (**Figure 2E**). Aligned with the increased abundance of PUFAs, the expression of *D6D* and *D5D* was upregulated upon early differentiation of ESCs (**Figure 2F**) and TSCs (**Figure 2G**).

**Figure 2.**
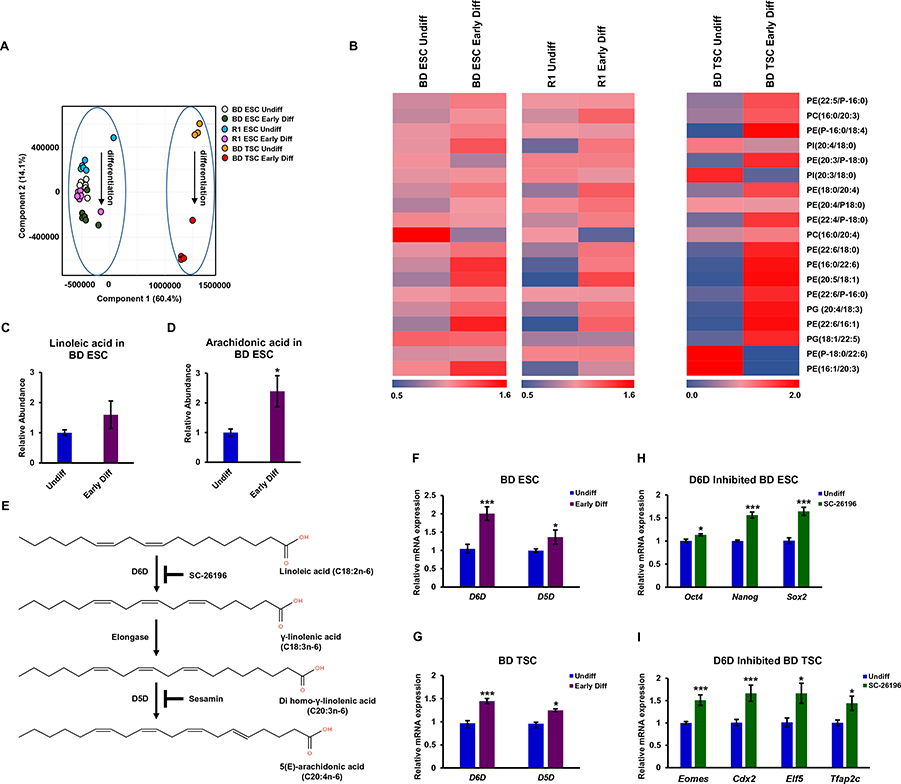
Polyunsaturated lipids are a metabolic signature of early differentiating ESCs and TSCs. ESCs and TSCs of the same background (129 / black 6) were seeded in Tx defined culture medium. After 24 hours, medium was changed to fresh medium with (“Undiff”) or without (“Early Diff”) factors that support self-renewal. Their metabolome was examined after 48 hours, using 0.5 × 10^6^ cells per sample. The second principal component (PC2) of a partial least squares discriminant analysis (PLS-DA; A) shows discrete metabolome of the differentiating SCs. R^2^ = 0.93, Q^2^ = 0.83 (n = 6). A heat map representation of the abundance of identified differential polyunsaturated phospholipids of the dataset shown in A, following variable importance in projection (VIP; > 1) analysis (B). The abundance of each metabolic feature was normalized to the sum intensity of peaks of each chromatogram. The colors of quadrants in the heat maps represent the relative abundance of the metabolites, after normalization to the mean abundance of each metabolite across all samples (n = 6). Following hydrolysis of lipids in ESC extracts, total abundance of linoleic acid (C) and arachidonic acid (D) was measured by GC-MS and normalized to the sum of peaks area. The biosynthesis of PUFAs starts from essential fatty acids with two double bonds, and is catalyzed by D6D and D5D (E). *D6D* and *D5D* expression is upregulated in early differentiating ESCs (F) and TSCs (G), as measured by RT-PCR (n = 6). ESCs and TSCs (129 / black 6 background) were treated by the D6D inhibitor SC-26196 (0.2 μM) or DMSO control for 48 hours in feeder free culture without 2i/LIF or FGF4/TGFβ/heparin and the expression of stem cell markers was measured by RT-qPCR (H-I; n = 4). Values are the mean ± SEM. *, P < 0.05; **, P < 0.01; ***, P < 0.001.

To examine a possible influence of D6D on the expression of markers of stemness in differentiation-inducing culture conditions, we used SC-26196, a specific inhibitor of D6D. D6D inhibition retained the expression of pluripotency marker genes in early differentiating ESCs (**Figure 2H)**, and TSC marker genes in differentiating TSCs (**Figure 2I)**.

Taken together, these results suggest that differentiated ESCs and TSCs share increased levels of polyunsaturated lipids. This shift can be attributed to D6D and D5D expression, and is allocated at the exit from a self-renewing state.

We wished to better define the influence of polydesaturation on blastocyst derived SCs. For this, we inhibited D6D or D5D in ESCs or TSCs of the same (129 / black 6) background (main Figures), and validated the data from these experiments using independent ESC and TSC lines (Supplemental Figures).

Microscopic examination of the cells suggested a higher number of colonies and greater colony size in D6D inhibited ESCs. Indeed, D6D or D5D inhibition caused an increase in ESC numbers (**Figure 3A-B)**, concomitant to an increase in clonogenicity (**Figure 3C-D)**. D6D inhibition in ESCs in a defined Tx medium also resulted in increased proliferation of ESCs (**Figure 3E)**. The effect of D6D inhibition on self-renewal was dose-dependent (**Figure 3F)**, and specific, as confirmed by the inhibition of D6D by SiRNA (**Figure 3G**).

**Figure 3.**
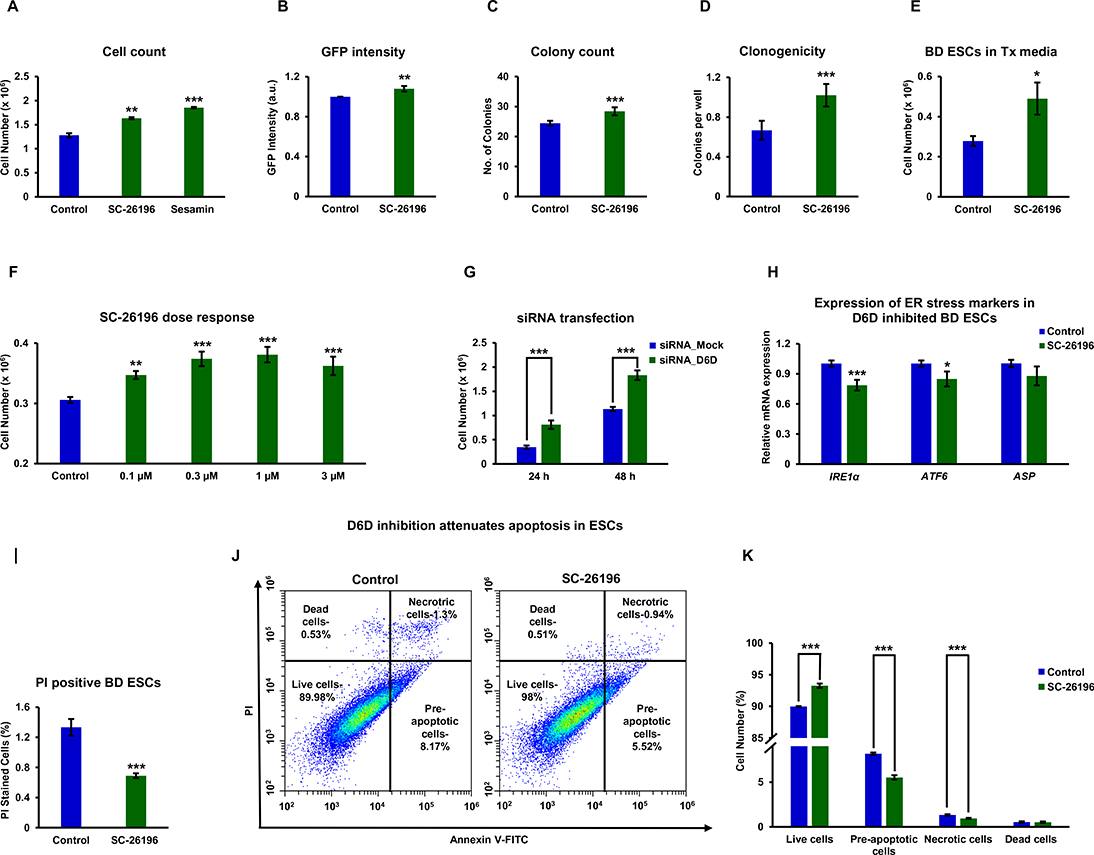
Polydesaturation of lipids causes ER stress-related apoptosis in ESCs. ESCs (129 / black 6 background in A-I; V6.5 in J-K) were treated with the specific D6D inhibitor SC-26196 (0.2 μM), D5D inhibitor sesamin (10 μM), or DMSO control for 48 hours in feeder free culture. Cell counting was carried out using CytoSmart cell counter and validated by manual counting (A; n = 4). Fluorescence was quantified in *Oct4*-GFP labeled 129 / black 6 ESCs using a plate reader to verify the increase in proliferation (B; n = 4). Colony numbers per field were estimated under the microscope (C; 5 fields taken for each well, for 4 wells). Cell survival was further evaluated by a clonogenicity assay (D; (n = 48). Cell count was further carried out in defined Tx medium (E; n = 4). Dose response evaluation of the influence of D6D inhibition on cell proliferation rate was performed with 0.1-3 μM SC-26196 (F; n = 4). For validation of the specific effect on D6D, siRNA was applied (G; n = 4). The expression of the ER stress activators *IRE1α*, *ATF6*, and *ASP* was assessed by RT-qPCR upon D6D inhibition for 24 hours (H; n = 4). The percentage of dead cells in D6D inhibited ESCs was further assessed by propidium iodide (PI) staining (I; n = 4). To evaluate apoptosis in V6.5 ESCs (as they have no GFP label) following 24 hours of D6D inhibition, we used a kit based on annexin V-FITC staining. A representative dot plot and the corresponding bar graph of one of four experiments (n = 4 for each) is presented (J-K). Data are presented as mean ± SEM. *, P < 0.05; **, P < 0.01; ***, P < 0.001.

The ability of cells to respond to perturbations in endoplasmic reticulum (ER) function is critical for cell survival (46). Alterations in ER lipid composition result in ER stress (47). In particular, ER function is associated with the degree of desaturation of cellular fatty acyl chains (48, 49). We assumed that polydesaturation may decrease the viability of ESCs by driving ER stress-related apoptosis. In line with this possibility, we found reduced expression of the ER stress activators inositol-requiring transmembrane kinase/endoribonuclease 1α (*IRE1*α) and activating transcription factor 6 (*ATF6*) upon D6D inhibition (**Figure 3H**). D6D inhibition reduced the percentage of dead (**Figure 3I**) and increased the percentage of pre-apoptotic and necrotic (**Figure 3J-K**) ESCs, confirming an anti-apoptotic effect. To further confirm the pro-apoptotic influence of D6D in ESCs, we tested its effect on the viability of further ESC lines (**Figure S1**). We also wished to define the effect of D6D inhibition on ESC cell cycle, and found no effect (**Figure S1**), suggesting that the higher proliferation rate is not due to cell cycle rate, and may be attributed to less ER stress-related apoptosis, leading to higher viability of ESCs following downregulation of D6D.

Similarly to ESCs, the inhibition of polydesaturation by D6D or D5D increased TSC proliferation (**Figure 4A**), and the number of colonies (**Figure 4B**), concomitantly with decreased expression of IRE1α (**Figure 4C**), and attenuation of apoptosis (**Figure 4D-E**). Unlike in ESCs, the inhibition of D6D also increased the percentage of TSCs in S phase (**Figure 4F-G**), suggesting that the increase in proliferation rate in TSCs is partially mediated by altered cell cycle kinetics. We validated the increased proliferation of TSCs using independent TSC lines (**Figure S2**).

**Figure 4.**
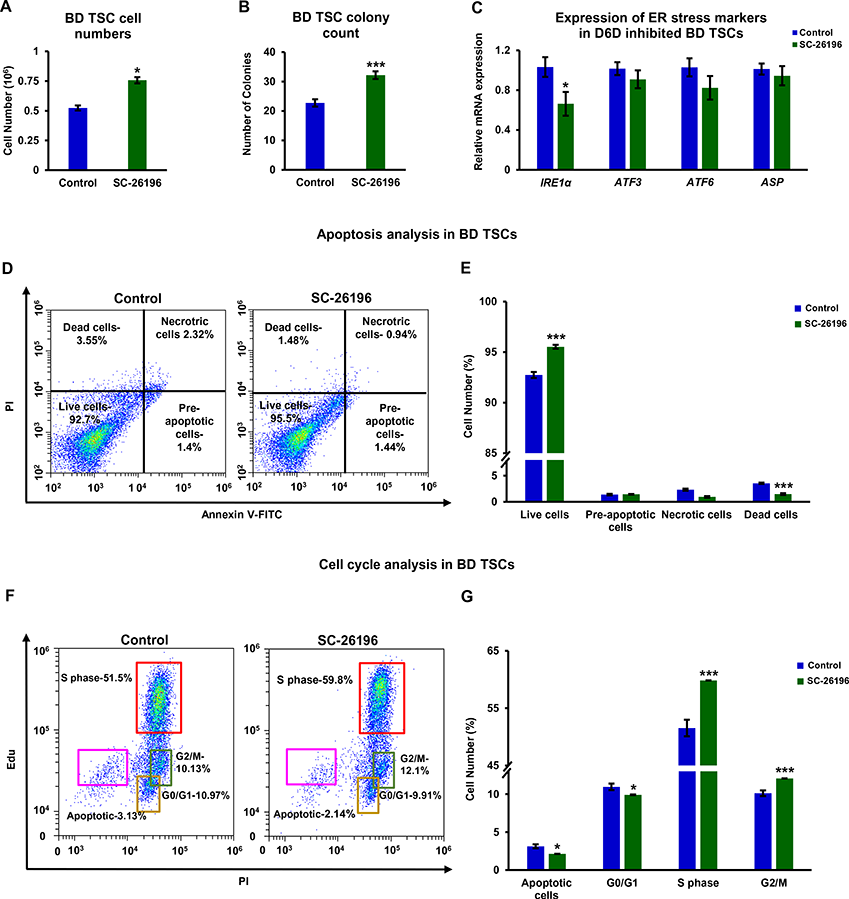
D6D causes ER stress-related apoptosis and modifies cell cycle in TSCs. TSCs (129 / black 6) were treated with SC-26196 (0.2 μM) or DMSO control for 48 hours in feeder free culture. Cell counting was carried out using CytoSmart cell counter and validated by manual counting (A; n = 4). Colony numbers per field were estimated under the microscope (B; 5 fields taken for each well, for 4 wells). The expression of *IRE1*, *ATF6, ATF3* and *ASP* was examined by RT-qPCR following D6D inhibition for 24 h (C; n = 4). Apoptosis was assessed in TSCs following D6D inhibition for 24 hours using Annexin V/PI staining (D-E; n = 4). The numbers in each quadrant of dot plot represent the percentage of cells. Cell cycle was analyzed following inhibition of D6D by SC-26196 for 24 hours. The numbers in each quadrant of dot plot represent the percentage of cells (F-G; n = 4). Data are presented as mean ± SEM. *, P < 0.05; ***, P < 0.001.

The cultivation of ESCs and TSCs is mostly done on feeder cells that release factors that support SC cultivation. To evaluate the influence of D6D on SC proliferation in feeder cell-based culture, we carried out FACS analyses of the ratio of SCs / mouse embryonic fibroblasts (MEFs) in culture. We used *Oct4*-GFP labeled ESCs for the quantification of ESCs. For the quantification of TSC proliferation in co-culture, we labeled TSCs using an anti-CD40 (a TSC marker) antibody. ESC relative proliferation in culture with feeder cells increased following the inhibition of D6D, whereas the number of D6D inhibited TSCs only slightly and insignificantly increased in culture with the MEF feeder cells (**Figure S3**).

A balance between mono- and poly-desaturation of lipids is maintained in the liver, brain, lymphocytes and adipocytes, as PUFAs represses Scd1 activity and mRNA stability (50–52). Scd1 inhibition activates the IRE1/XBP1s signaling arm of the unfolded protein response, causing ER stress (53, 54). Interestingly, the stress response following Scd1 inhibition causes massive apoptosis in human ESCs, but not somatic cells; mouse ESCs showed partial elimination (37). Accordingly, we reasoned that the increase in cell viability in D6D inhibited ESCs may result from upregulation of Scd1 activity, leading to lower apoptosis rates. In line with this hypothesis, *Scd1* was upregulated in ESCs following inhibition of D6D (**Figure 5A**). Surprisingly, in contrast to the inverse regulation of *Scd1* by D6D in ESCs, *Scd1* was downregulated upon D6D inhibition in TSCs, suggesting a reciprocal regulation of Scd1 in the two cell types by PUFAs (**Figure 5B**). Scd1 mRNA expression was slightly downregulated in early differentiating ESCs, whereas a non-significant upregulation noted in TSCs (**Figure 5C**). We wished to further examine possible functional implications of Scd1 expression in blastocyst-derived SCs. The expression of the pluripotent markers *Oct4* and *Sox2* was downregulated in early differentiating ESCs (**Figure 5D**), whereas *cdx2* and *Tfap2c* were upregulated in early differentiating TSCs (**Figure 5E**) following inhibition of Scd1. The inhibition of Scd1 by PluriSIn 1 was reported by Ben-David and colleagues to partially eliminate mouse ESCs (37). Aligned with the results of this study, we saw partial elimination of mouse ESCs by PluriSIn 1 or A-939572 - another Scd1 specific inhibitor (**Figure 5F-G**). Remarkably, the inhibition of Scd1 in TSCs increased their viability (**Figure 5H-I**). The addition of conjugated oleate to Scd1-inhibited TSCs attenuated their proliferation, reducing it to the level seen in control cells (**Figure 5J**). As expected from the higher viability of TSCs upon Scd1 inhibition, they showed decrease in the expression of ER stress genes **(Figure 5K)**, and reduced apoptosis **(Figure 5L-M)**. We tested the viability of ESCs and TSCs under Scd1 inhibition and confirmed that the increase in viability of TSCs, whereas decrease of ESC viability by Scd1 inhibition is not cell line specific (**Figure S4**). Inhibition of Scd1 in feeder cell-based culture resulted in increased proliferation in TSCs, whereas lower ESC proliferation (**Figure S5**).

**Figure 5.**
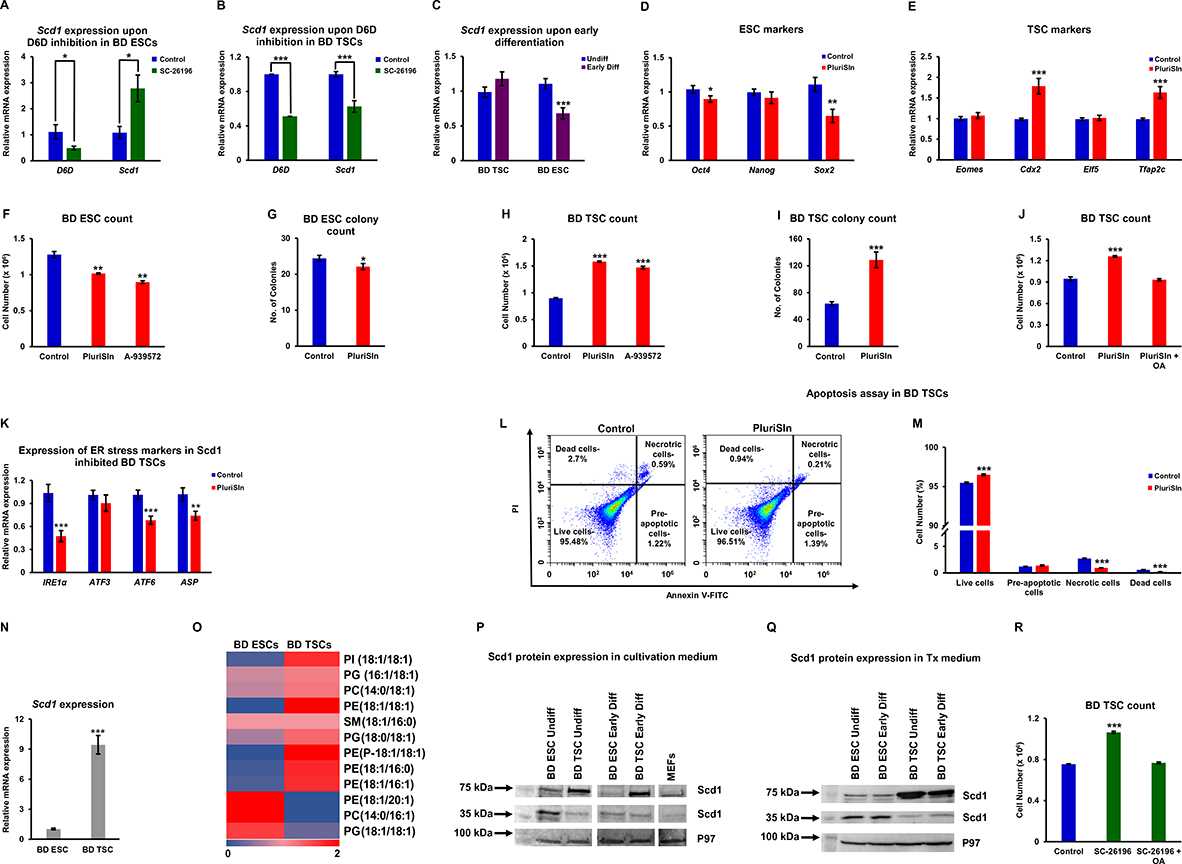
D6D regulates Scd1 expression, essential for ESCs, but deleterious for TSCs. The expression of *Scd1* in feeder free culture following 48 hours of inhibition of D6D (0.2 μM SC-26196), or in DMSO-treated control cells, was evaluated by RT-qPCR in undifferentiated ESCs (A) and TSCs (B) (both 129 / black 6 background). *Scd1* expression was also assessed in early differentiating ESCs and TSCs, compared to their undifferentiated counterparts (C). The influence of Scd1 on pluripotency (D) or TSC (E) markers in early differentiating SCs, as well as on cell viability (F-J) was evaluated following treatment with Scd1 inhibitors [PluriSIn 1 (20 μM) or A939571 (300 nM)] or DMSO vehicle for 48 hours. Conjugated oleate (100 μM) was added to PluriSin 1-treated TSC culture (J). The expression of *IRE1α*, *ATF3*,*ATF6*, and *ASP* was assessed by RT-qPCR following inhibition of Scd1 by PluriSIn 1 for 48 hours (K), and concomitant apoptosis was evaluated using an Annexin V-FITC-based kit (L-M). The relative expression of *Scd1* in self-renewing ESCs and TSCs was evaluated by RT-qPCR (N). Monounsaturated fatty acyl chain phospholipids were identified and quantified in self-renewing TSCs and ESCs by LC-MS. The abundance of each metabolic feature was normalized to the sum intensity of peaks of each chromatogram. The colors of quadrants in the heat maps represent the relative abundance of the metabolites, after normalization to the mean abundance of each metabolite across all samples (n = 6; O). Scd1 protein expression was determined by Western blot analysis (10% acrylamide gel) in self-renewing ESCs and TSCs in ESC cultivation medium or Tx medium, respectively (P), or both cell types in the same defined Tx medium (Q). TSC proliferation was evaluated in TSCs, as above, following incubation with SC-26196 or SC-26196+conjugated oleate (100 μM) (n = 4; R). Data are presented as mean ± SEM. n=4. *, P < 0.05; **, P < 0.01; ***, P < 0.001.

We assumed that the detrimental influence of Scd1 on ESCs may reflect a differential expression of the enzyme and its products. We found approximately one order of magnitude higher gene expression of *Scd1* in TSCs (**Figure 5N**), and the expected higher accumulation of MUFAs (**Figure 5O**). The examination of Scd1 protein expression suggested a surprising alteration in of Scd1 in ESCs (molecular mass ~35 kDa) and TSCs (molecular mass ~70 kDa) (**Figure 5P; Figure S6**). Upon early differentiation, Scd1 protein expression is decreased in ESCs, with no change noted in early differentiating TSCs (**Figure 5P; Figure S6**). MEFs (differentiated embryonic cells), show no preference to either of the Scd1 variants (**Figure 5P; Figure S6**). Scd1 is regulated by the availability of fatty acids (50). To better understand the shift in Scd1 variants and address a possible regulation of their expression by the availability of external fatty acids, we incubated ESCs in 0-10% FBS (as a source of external fatty acids). Although we noted an upregulation of Scd1 with increasing percentage of serum, no change in the high molecular mass variant of Scd1 was noted (**Figure S6**). Nevertheless, we then examined the expression of Scd1 in ESCs and TSCs, cultivated in the same defined (Tx) medium, and saw the same alteration in the Scd1 variant (**Figure 5Q**). Notably, oleic acid – the product of Scd1, blocked the Scd1-dependent increase in viability in D6D inhibited TSCs (**Figure 5R**), suggesting that this increase is mediated by the downregulation of the basal high MUFA levels in TSCs.

## DISCUSSION

In the current study, we characterized the metabolic profiles of early differentiating ESCs and extra-embryonic TSCs vis-à-vis their self-renewing counterparts. Our metabolomics analyses clearly show a metabolic shift in both blastocyst-derived SCs as they exit self-renewal. This shift results in higher levels of PUFAs, following increased expression of D6D and D5D. No change was seen in the levels of linoleic acid. The inhibition of lipid polydesaturation by D6D or D5D retains self-renewal in ESCs and TSCs, suggesting that PUFA levels are driving differentiation in both cell types, and that controlling lipid desaturation is essential for the balance between self-renewal and differentiation and for the maintenance of ESCs and TSCs in culture.

Previous work suggests that the global metabolome of ESCs is characterized by abundant metabolites with highly unsaturated structures, of which the levels decrease upon differentiation (9), whereas the results of our analyses demonstrate upregulation of lipid desaturation upon differentiation. This seeming discrepancy may be a result of our focus on desaturation of lipids. It may also be attributed to variations in the differentiation stage studied (early differentiated vs fully differentiated neurons or cardiomyocytes). Despite this seeming discrepancy, our data on the retention of pluripotency by the inhibition of lipid polydesaturation in ESCs are well aligned with the results of Yanes and colleagues on ESC differentiation (9). Our results further link the shift in lipid polydesaturation to lower SC self-renewal by exerting ER stress-related apoptosis, and to the early differentiation of extra-embryonic blastocyst-derived SCs.

The regulation of the ER status by lipid desaturation has been reported in multiple studies in somatic cells. In ESCs, downregulation of Scd1 causes ER stress and apoptosis, as demonstrated by Ben-David and colleagues (37), and by our results. Given the regulation of Scd1 by PUFAs in the liver, lymphocyte, brain, and adipocytes cells (50), we reasoned that lipid polydesaturation inversely regulates Scd1 in ESCs. Our results are in concurrence with this assumption, suggesting that polydesaturation may modulate the ER status via regulation of Scd1 expression.

Surprisingly, we saw a downregulation of Scd1 following D6D inhibition in TSCs. Despite the reciprocal regulation of Scd1 by D6D in ESCs vs TSCs, inhibition of polydesaturation decreases ER stress and apoptosis in both cell types. The fact that Scd1 inhibition increases ESC death while increasing the viability of TSCs provides an explanation for this, and underscores a unique essentiality of MUFAs in ESCs when compared to their extra-embryonic counterparts.

The findings that TSCs express higher mRNA levels and a different protein variant of Scd1, provide mechanistic insights into the differences in its essentiality and function. A high molecular Scd1 variant, such as the one expressed in TSCs, was previously attributed to the formation of dimers (55, 56). It was suggested to have potentially altered function or stability (57), but it has not been isolated, and its function has so far not been studied .

Our results show that while ESCs are addicted to MUFAs, TSCs have high levels of MUFAs, high mRNA expression of Scd1, and an altered variant of Scd1 protein. The addition of conjugated oleate to D6D-inhibited TSCs rescues apoptosis, supporting the notion that increasing MUFA levels may be detrimental for TSCs, and linking the pro-apoptotic effect of D6D to the regulation of MUFA levels.

Further research is required to establish the function of Scd1 variants and their regulation by PUFAs in the early embryo. Seeing that a high molecular variant of Scd1 is expressed in the extraembryonic TSCs, it would be interesting to study its expression and function in the placenta.

The downregulation of the expression of pluripotency markers by Scd1 inhibition in ESCs may be mediated by a non-ER influence of the enzyme. The upregulation of TSC marker genes by Scd1 inhibition suggests that the regulation of the blastocyst-derived SC populations by Scd1 is also mediated by modulation of the potency of the cells to differentiate, and further underscores its pivotal role in the regulation of the two cell populations and their progenies.

ESCs or TSCs are mostly cultivated in co-cultures with feeder cells that provide them with nutrients and paracrine signals required for their cultivation. Our analyses show that the regulation of ESC viability by lipid desaturases is less pronounced in co-culture with MEFs than in feeder free culture, but still significant. This is likely due to the differences in the media lipid composition, conferred by the feeder cells. We saw no effect of D6D inhibition in TSCs in co-culture, implying that the advantage offered by inhibition of polydesaturation is limited to their cultivation in feeder free culture.

Taken together, our data suggest that lipid polydesaturation is a common metabolic signature of differentiating blastocyst-derived SCs, marking a shift from self-renewal to differentiation. This shift is pro-apoptotic in both ESCs and TSCs via downregulation of Scd1 in ESCs (dominant variant~35 kDa), whereas upregulation of Scd1 (showing a ~70 kDa variant) in TSCs. Our results expand the concept of metabolic regulation of stem cells fate decisions to the extra-embryonic stem cells. Our work on blastocyst-derived cells suggests that the availability of MUFAs and PUFAs in the microenvironment of the blastocyst, and the expression of D6D, D5D and Scd1 in the blastocyst cell types are critical regulators of the concerted differentiation and expansion of the blastocyst cell populations. Such premise requires further studies, and especially in mammalian embryos.

## MATERIALS AND METHODS

### Cell lines

R1 *Oct4*-GFP ESCs were kindly provided by Prof. Andras Nagy of the Lunenfeld Research Institute. V6.5 ESCs were kindly provided by Rudolf Jaenisch, MIT. TNGA Nanog ESCs (with a background of 129 / black 6) and the blastocyst-derived TSC clones: BD TSC 1 with a background of 129 / black 6, and BD TSC 2 with Homo BL6 background were kindly provided by Dr. Yosef Buganim, Hebrew University of Jerusalem.

### Cell culture

Cells were cultured in standard conditions. In brief: Cells were maintained in a humidified incubator at 5% CO_2_ and 37 °C (Thermo Fisher Scientific, USA).

ESCs in maintenance were cultured as previously described by us (17, 58), on a confluent adhesive layer of mitomycin-inactivated MEF feeder cells in 2i ESC culture DMEM medium supplemented with 15% Fetal bovine serum (FBS; Gibco, USA), 1.5% Pen/Strep (Biological Industries, USA), 1% sodium pyruvate (Biological Industries, USA), 1% L-glutamine (Biological Industries, USA), 1% non-essential amino acids (Gibco, USA), 0.1 mM 2-mercaptoethanol (Gibco, USA), 2.8 μM CHIR99021 (Biogems, Peprotech), 1 μM PD0325901 (Biogems, Peprotech), mouse Leukemia inhibitory factor (LIF; in-house production). Every two days, cells were passaged. Before experiments, ESCs were cultured in feeder free conditions (in LIF/2i ESC culture medium) on 0.2% gelatin coated culture plates for 3 passages.

TSCs in maintenance were cultivated as previously described (4), on gelatin coated mitomycin inactivated MEFs in 30% TSC medium [RPMI 1640 (Gibco, USA) supplemented with 20% FBS (Gibco, USA), 1% non-essential amino acids (Gibco, USA), 1% L-glutamine (Biological Industries, USA), 1% sodium pyruvate (Biological Industries, USA), 1% pen/strep (Biological Industries, USA), 50 μM β-mercaptoethanol (Gibco, USA)] and 70% MEF conditioned medium (medium collected from mitomycin inactivated MEFs, containing the same supplements) freshly supplemented with 25 ng/mL of human recombinant FGF4 (prepared in house) and 1 μg/mL heparin (Sigma-Aldrich, USA). TSCs were passaged every 3-5 days, using trypsin EDTA. For feeder free experiments, TSCs were cultured in Matrigel ® Matrix (Corning Life Sciences, USA) coated [1:30 dilution in DMEM/F-12 medium (Gibco, USA)] tissue culture plates in Tx defined medium (59) with freshly added 25 ng/mL of mouse recombinant FGF4 (In-house production), 1 μg/mL heparin (Sigma-Aldrich, USA), 1 μg/mL of human TGFβ1 (Biogems, Peprotech).

Fetal bovine serum used throughout the study was from one batch, to avoid possible batch-dependent changes in the lipid composition of the cultivation medium.

### ESC and TSC early differentiation

Cells were cultivated in a feeder free medium: ESCs in 2i culture medium on 0.2% gelatin coated plates and TSCs on Matrigel coated plates in Tx defined medium (59). Cells were maintained in feeder free conditions for at least 3 passages prior to experiments. For initiation of differentiation, growth factors and cytokines that maintain cells in pluripotency (LIF and 2i) or TSC state (heparin, recombinant human TGF-β and FGF-4) were excluded from medium. For the infliction of differentiation, 24 hours following seeding, we allowed SCs to differentiate by excluding the above factors from cultivation medium for 48 hours. The medium of SCs of the same experimental lot was replaced with fresh ESC/TSC culture medium as above, and they were used as undifferentiated control. We then compared the metabolic composition of early differentiating SCs to that of undifferentiated cells.

### Metabolomics analyses

Following 0-48 hours of spontaneous differentiation, ESCs or TSCs were harvested and centrifuged at 200 g, 4 °C, 5 min. Cell pellets were washed twice with Dulbecco’s PBS without Ca and Mg (Biological Industries, USA), snap frozen in liquid nitrogen, and kept at −80 °C. Cell pellets were extracted in an ice cold solvent containing 86.5% methanol (J. T. Baker, USA), 12.5% LC-MS grade water (J. T. Baker, USA), 1% formic acid (Tokyo Chemical Industry Co., Ltd., Japan). Samples were vortexed and extracted by ultrasonication followed by freeze-thaw cycles. Ultrasonication was carried out using Bioruptor® Plus sonicator (Diagenode, USA) at 30 second on / off pulsation cycles for 5 min. Following centrifugation, ultra-sonication and freezing cycles were repeated for two additional cycles and pooled for each sample in a fresh pre-labeled glass tube. Supernatant was transferred to clean glass tubes, and concentrated in a SC210A speed vacuum concentrator (SAVANT SPD 121P, Thermo scientific, USA) at 30 °C. The remaining pellet was extracted in 400 μL acetone, ultra-sonicated for 30 seconds, and centrifuged for 10 min at 4 °C. Supernatant was transferred to the respective glass tubes containing the methanol/water concentrated extracts and concentrated to dryness. Samples were reconstituted in 200 μL of reconstitution solvent: 95% acetonitrile (J. T Baker, USA; LC-MS grade), 5% water (LC-MS grade; J. T Baker, USA), 0.1% formic acid (Tokyo Chemical Industry Co., Ltd., Japan), and transferred to 96 well 700 μl sample plate (Waters^®^Acquity UPLC plate, USA).

Ten microliters of samples were injected to a Waters Acquity UPLC H-Class apparatus (Waters, Milford, MA, USA) equipped with UPLC CSH C18 2.1 x100 mm, 1.7 μm column (Waters, Ireland). The column was equilibrated with 0.4 mL/min flow of 40% of mobile phase A (ACN: Water 2:98% V/V) and 60% of mobile phase B (ACN). The linear gradient program was as follows: 60% mobile phase A (0.1% formic acid in water) and 40% mobile phase B for 1 min. The mobile phase B proportion was increased to 70% (v/v) in 5 min. From 5 to 8 min, mobile phase consisted of 24% A, 40% B, and 36% C [(isopropanol (Chemsolute, T.Geyer)]; from 8 to 9 min, 20% A, 35% B, and 45% C; from 9 to 12 min, 18.4% A, 33% B, and 48.6% C; from 12 to 17 min, 12% A, 25% B, and 63% C; and up to 25 min, 0.4% A, 10.5% B, and 89.1% C. Aquity UPLC system was coupled to a Xevo G2-XS, high resolution and high mass accuracy QTOF equipped with ESI. Negative ion mode was used for downstream analyses. Masses of 30-2000 Dalton underwent MS^E^ analysis with ramp collision energies 30-60eV. Mass range calibration was performed using Leucine enkephalin (Waters, USA) infused every 0.3 seconds. A quality control (QC) mix prepared from pooled samples was injected every 10 runs. A blank sample was injected every 10 samples. A minimum intensity cutoff mass was set to 100 m/z. Fold change greater than 100 from blank and lowest mean abundance in blank was set as another threshold. Data acquisition and visualization was performed using MassLynx 4.1 (Waters Co., UK). Spectral deconvolution, alignment, and normalization were performed using Progenesis QI (Nonlinear dynamics, Waters Corporation, UK). For metabolite identification, exact mass, isotope patterns and fragmentation patterns were compared to 18 metabolite libraries compatible with Progenesis QI (60). Multivariate statistical analyses, masses were carried out using Progenesis QI and Metaboanalyst 4.0 (Waters Corporation, UK). Feature intensities were quantile normalized.

### Evaluation of total arachidonic acid and linoleic acid levels using GC-MS

ESCs or TSCs were harvested and washed twice with PBS. Cell pellet (1×10^6^ cells) was snap frozen in liquid nitrogen and lyophilized (Labconco Corporation, USA). Lyophilized cell pellets were hydrolyzed with 1 mL of 1M KOH (Sigma Aldrich, USA) in 70% Ethanol (JT Baker, USA; HPLC grade) at 90 °C for 1 hour. The reaction mixture was acidified with 0.2 mL of 6M HCl (Sigma Aldrich, USA) and then 1 mL of water was added. The hydrolyzed fatty acids were extracted with 1 mL of hexane (GC grade, Sigma Aldrich, USA) and further concentrated using vacuum concentrator (Savant SPD121P speedVac concentrator, Thermo Scientific, USA). Fatty acids were then methylated with 2 mL of Boron Trifluoride (BF3) in Methanol (14% W/V) (Sigma Aldrich, USA) and incubated at 55 °C for 1.5 hours with intermittent shaking every 20 min. 2 mL of saturated solution of NaHCO_3_ (Sigma Aldrich, USA) and 3 mL of hexane were then added, and the tubes were vortex-mixed for a minute. The hexane layer was recovered carefully and concentrated using vacuum concentrator. Samples were reconstituted in 100 μL hexane (GC grade, Sigma Aldrich, USA). GC-MS analysis was performed using an Agilent 8860 GC system equipped with an Agilent 7683 N auto sampler and a Agilent J & W GC columns (30 m × 0.25 mm × 0.2 μm, Agilent technologies USA). The injector was set at 220 °C and 1 μL injections were made with helium as carrier gas, maintained at flow rate of 1 mL/min.

GC temperature gradient is given as **Table S2**. The transfer line from GC column to MS (Agilent 5977B GC/MSD) was set to 250 °C, ionization energy of 70 eV. Standards for arachidonic acid and linoleic acid (Sigma Aldrich, USA) were used for validation of identification.

### Scd1 or D6D enzyme inhibition assay

Selective inhibitors for Scd1 PluriSIn-1 and A-939572, D6D (SC-26196) and D5D (sesamin) were purchased from Cayman Chemicals, USA. The effect of the pharmacological inhibitors was evaluated for ESCs and TSCs following 48 hours incubation. Oleate conjugated to albumin (Sigma Aldrich, USA) was also tested for its influence on Scd1 or D6D-inhibited TSCs.

### SiRNA transfection

ESCs were maintained at a density of 2.5 × 10^4^ cells/well in 24 well plate for 24 hours following passage, in ESC LIF/2i medium. Cells were transfected with MISSION® SiRNA oligomers or MISSION® siRNA Universal Negative Control #1 (Sigma Aldrich, Merck, USA) at a final concentration 20 nM, mixed with 1.5 μL X-tremeGENE transfection reagent (Sigma Aldrich, Merck, USA). The complex was incubated at room temperature for 30 min in serum free medium and transferred to the cells. Following 6 hours post transfection, the medium was replaced with ESC LIF/2i medium, and the cells were harvested after 48 hours.

### Cell counts

Cells were seeded at a density of 15,000 (ESCs) or 50,000 (TSCs) cells per well in 24 well plates, in feeder free culture. Cell count was carried out using Cyto Smart cell counter (Corning Incorporated Life Sciences, USA). Counts were validated by manual counting.

### Fluorescent Detection

Fluorescence of *Oct4*-GFP ESCs was further quantified using CYTATION 3 plate reader (LUMITRON, Israel) at an excitation wavelength of 395 nm and emission peak of 509 nm.

### Microscopy

Microscopic imaging was taken at 10 × magnification using OLYMPUS IX73 Inverted Microscope (OLYMPUS Life Science, USA).

### Quantitative real-time Polymerase Chain Reaction (RT qPCR)

Total RNA for the experiments was extracted from the cells using GENEzol™ TriRNA Pure Kit + DNAse (Geneaid HyLabs, Taiwan) according to the manufacturer’s protocol. Quantification of total RNA concentration was done using NanoDrop ND-1000 spectrophotometer (Version 3.8.1, Thermo Fisher). First strand cDNA synthesis was done from 1000 ng of RNA using qScript cDNA Synthesis kit (Quantabio, USA). RT qPCR was performed using Fast SYBR™ Green Master Mix (Applied Biosystems™, Thermo Fisher) and a CFX Connect™ Real-Time PCR Detection System. Primer sequences (HyLabs and IDT) are listed in **Table S3**. Relative expression of the transcripts was normalized to Hypoxanthine Phosphoribosyltransferase (HPRT)1 or Ubiquitin C (UBC), and the gene expression was quantified based on comparative 2^−ΔΔ*CT*^.

### Flow cytometry

Following dissociation, cells were centrifuged at 200 g for 5 min, and cell pellets were washed twice in PBS containing 1% FBS, and filtered through a cell strainer to remove aggregates. Cells incubated with secondary and no primary antibody (for stained cells) or unlabeled cells (for GFP-labeled cells) were used as negative controls. Flow cytometry was carried out on CytoFLEX Flow Cytometer (BECKMAN COULTER, Life Sciences) using a 100 μm nozzle. Analysis was carried out using CytExpert 2.4 software (BECKMAN COULTER, Life Sciences). For TSCs in co-culture with feeder cells, cells were incubated with anti-CD40 antibody (Ab22469, 1:300; Abcam), and then with Alexa Flour-conjugated anti Rat 488 secondary antibody (Ab150153, 1:500; Abcam).

### Apoptosis assay

Cells were seeded on 0.2% gelatin (ESCs) or matrigel (1:30 dilution; TSCs) coated culture plates. Cell pellets were washed with ice cold PBS and resuspended in 1X Annexin V binding buffer (Cell Signaling Technology, USA). Cells were then incubated with Annexin V-FITC conjugate (Cell Signaling Technology, USA) for 10 min on ice protected from light. Propidium Iodide (20 μg/mL; Fisher Scientific, USA) was added to cell suspension immediately before FACS analysis.

### Immunoblotting

Cells were collected by trypsinization, then washed with PBS (Biological Industries, Israel). Proteins were extracted with 100 μL in-house made RIPA (1% NP-40, 0.1%SDS, 1M Tris-HCL pH 7.4, NaCl, Sodium deoxycholate, EDTA, and water) + Protease Inhibitors (Roche, USA) and quantified using Bradford reagents (Sigma Aldrich, USA). Extracted proteins were reconstituted in an in-house Protein Sample Buffer (Glycerol 100%, 1M Tris pH 6.8, 10% SDS, β-mercaptoethanol, bromophenol blue) and heated for 10 min at 95 °C. Proteins were separated on 10% SDS-PAGE gels and transferred to 0.22 μm PVDF membrane (FroggaBio, USA) by an electric current at 300 mA for 70 min. The membrane was blocked in 5% BSA/TBST for 2 hours, then incubated with primary antibody overnight. Scd1 antibody was purchased from Cell Signaling Technology USA, (#2794, 1:1,000). Membrane was washed, then incubated with HRP conjugated secondary antibody (Abcam, USA, Ab 6721, 1:10,000). eIF4G2/p97 antibody from Cell Signaling Technology (#2182, 1: 3000) were used as a loading control. Chemiluminescence was detected by ECL (Advansta, USA), using Molecular Imager ChemiDoc™ XRS (Bio-Rad, USA).

### Cell cycle analysis

Cells were seeded at a density of 30,000 (ESCs) or 100,000 (TSCs) cells per well in a 6 well culture plate and treated with D6D inhibitor or DMSO vehicle for 24 hours. Upon 70% cell confluence, cells were treated with 10 μM of Edu (5-ethynyl-2′-deoxyuridine; Invitrogen™, Thermo Fisher Scientific, USA), and incubated for 30 min. After labeling, cells were harvested and washed twice with PBS. Cell pellets were resuspended in cold PBS and fixed by adding cold absolute ethanol dropwise while mixing. Cells were incubated overnight at 4 °C. Following centrifugation, cell pellets were permeabilized by adding 0.1% Triton® X-100 (Sigma Aldrich, USA) in PBS [1% BSA (Sigma Aldrich, USA)], and stained with Click-iT reaction mix [50 mM Trizma® Hydrochloride Buffer pH 7.5 (Sigma Aldrich, USA), 150 mM NaCL (Sigma Aldrich, USA), 2 mM CuSO4 (Sigma Aldrich), 2.5 μM Alexa Fluor 647 azide triethyl ammonium salt in DMSO (Thermo Fisher Scientific, USA) and 10 mM Sodium ascorbate (Sigma Aldrich, USA)] for 30 min. The cells were again washed with 0.1% Triton® X-100, 1% BSA in PBS, and stained with propidium iodide (Thermo Fisher Scientific, USA), 50 μg/mL of RNase (Thermo Fisher Scientific, USA) in PBS. EdU-analysis was performed using CytoFLEX Flow Cytometer and CytExpert 2.4 software (BECKMAN COULTER, Life Sciences).

## ACKNOWLEDGEMENTS

We thank Dr. Yosef (Yossi) Buganim for the kind gift of blastocyst isolated TSCs and ESCs and for providing the knowhow of TSC maintenance, and Avraham Greenberg and Prof. Itamar Simon for their kind help with the cell cycle analysis. We also thank Yosef Bar David for the help in using GC-MS instrument.

## AUTHOR CONTRIBUTIONS

A.M. conceived the project, supervised the project, analyzed data, secured funding, and wrote the manuscript. C.T.M. and R.B.H. maintained cell cultures and performed experiments, A.S. maintained cell cultures, P.O. performed experiments. A.D. assisted in preparing samples for analyses. A.Z., D.S., N.M.K., R.S. and S.A. analyzed data, I.P. validated the effect of D6D inhibition on a second line of ESCs. A.M. and C.T.M. prepared the manuscript for submission.

## ETHICS APPROVAL AND CONSENT TO PARTICIPATE

This study did not require ethical approval.

## DATA AVAILABILITY STATEMENT

The datasets used and/or analyzed during the current study are available from the corresponding author on reasonable request.

