## Supplementary file for "Lipid Desaturation Regulates the Balance between Self-renewal and Differentiation in Mouse Blastocyst-derived Stem Cells"

**Supplementary Information**

The following file contains supplementary material for the paper "**Lipid Desaturation Regulates the Balance between Self-renewal and Differentiation in Mouse Blastocyst-derived Stem Cells**".

This file is composed of:

- Supplementary figures and relative supplementary figure legends (6 figures)
- Uncropped Western blot pictures
- Supplementary tables (3 tables)

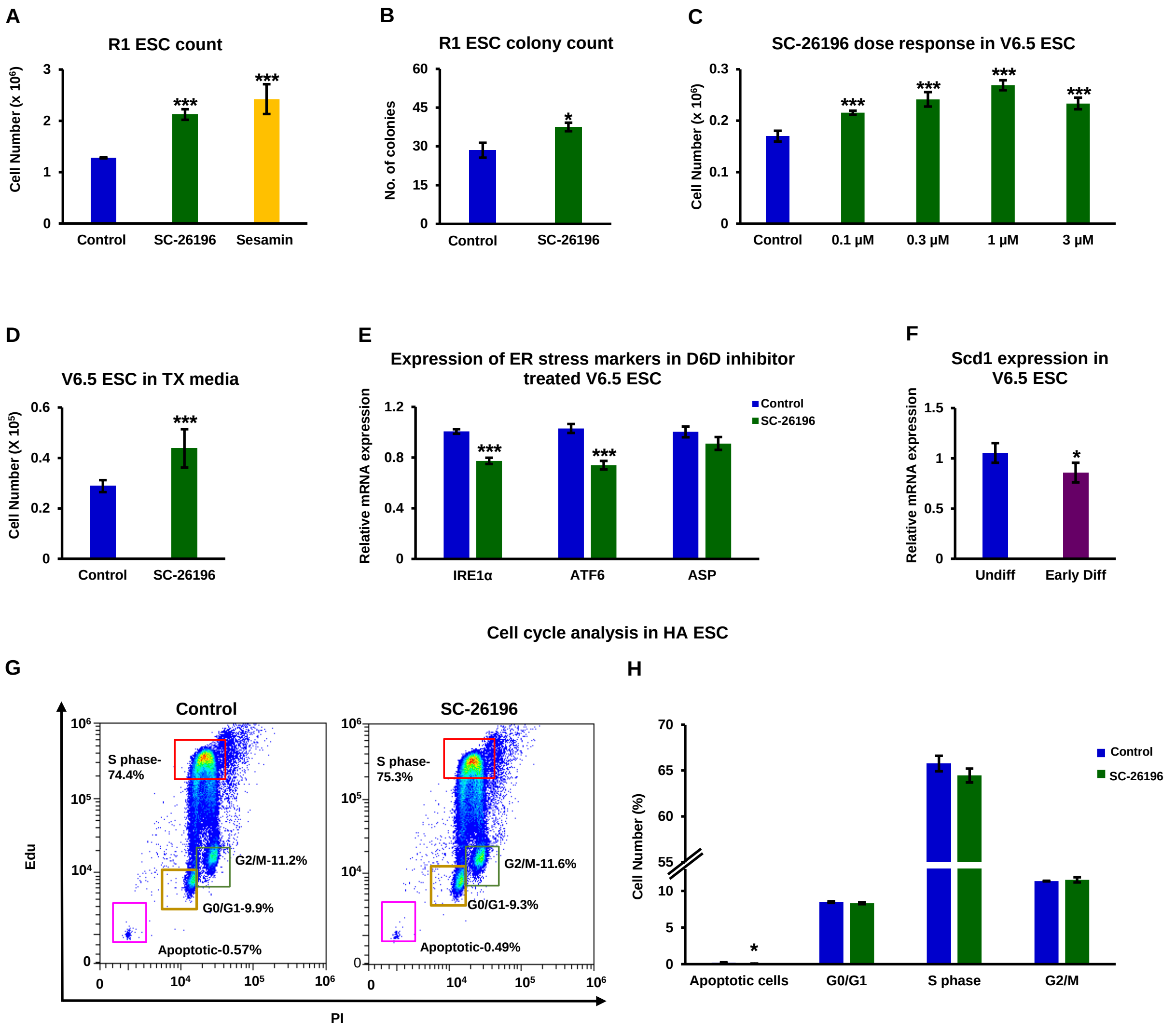


**Figure S1. ER stress-related apoptosis in ESCs due to polyunsaturation of lipids is not cell line specific, and not dependent on cell cycle.** To confirm the non-cell-line-specific nature of our results, we performed experiments with further ESC lines (R1 and V6.5; A-F). We also studied the effect of D6D inhibition on the ESC cell cycle (using the HA cell line used for the experiments shown in the main Figures), as an optional explanation for the increased proliferation rate. ESCs were treated with the specific D6D inhibitor SC-26196 (0.2 µM) or the specific D5D inhibitor sesamin (10 µM), or DMSO vehicle, for 48 hour in feeder free culture. Cell counting was carried out using CytoSmart cell counter and validated by manual counting (A; n = 4). Colony numbers per field were estimated under the microscope (B; 5 fields taken for each well, for 4 wells). Dose response evaluation of the influence of D6D inhibition on cell proliferation rate was performed with 0.1-3 µM SC-26196 (C; n = 4). The viability of ESCs was also assessed in Tx medium, as in A

(D; n = 4). The expression of ER stress markers was measured by RT-qPCR (E; n = 4). *Scd1* expression in early differentiating V6.5 ESCs was measured by RT-qPCR (F; n = 4). Cell cycle was analyzed in HA ESCs following inhibition of D6D by SC-26196 for 24 hours (G; n = 4). Data are presented as mean ± SEM. *, $P < 0.05$; **, $P < 0.01$; ***, $P < 0.001$.

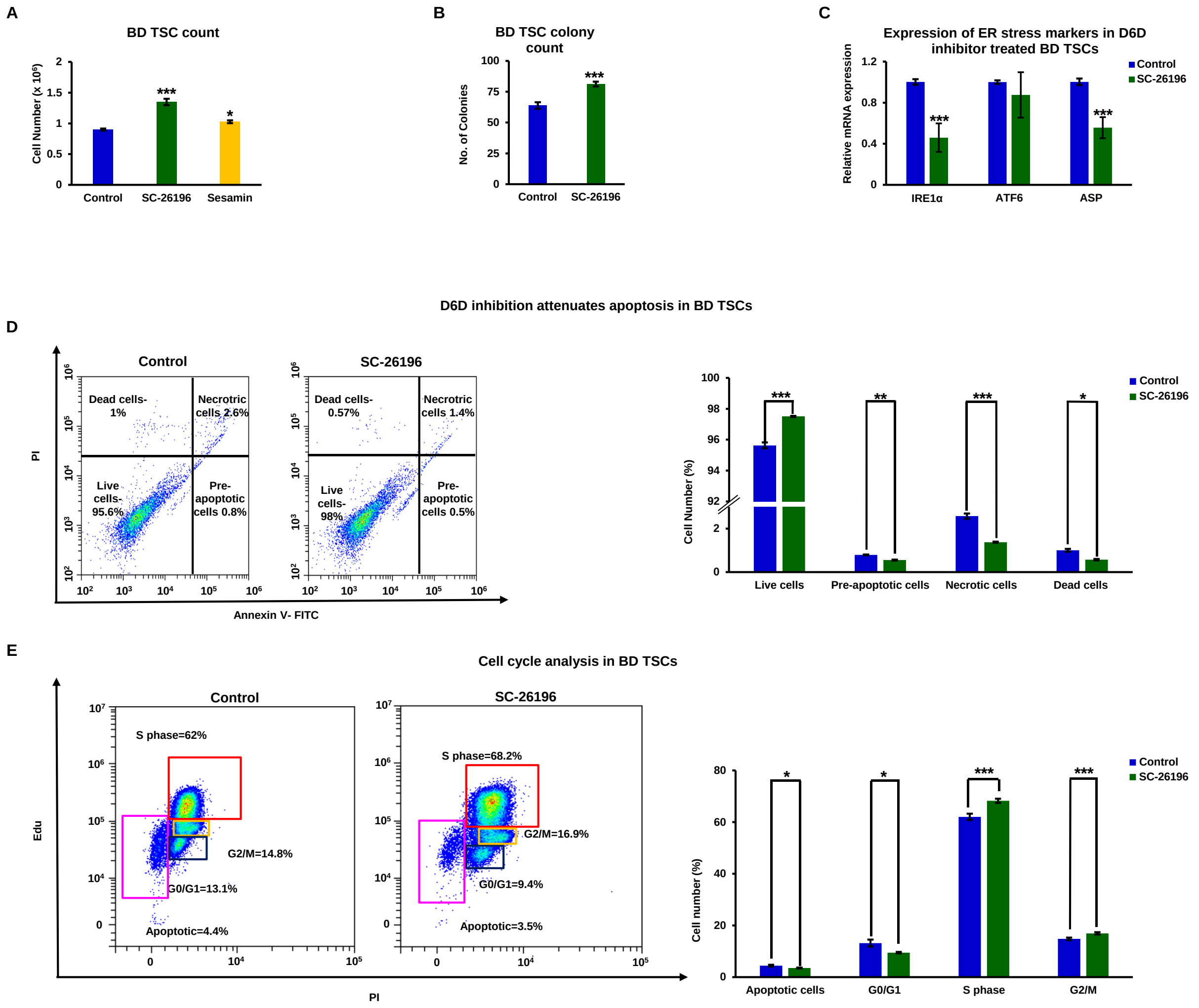


**Figure S2. ER stress-related apoptosis and changes in cell cycle rate in TSCs due to polyunsaturation of lipids are not cell line specific.** To confirm the non-cell-line-specific nature of our results, we performed experiments with a second blastocyst derived TSC line (Homo BL6 background). TSCs were treated with the specific D6D inhibitor SC-26196 (0.2 µM) or the specific D5D inhibitor sesamin (10 µM) (A-E) or DMSO vehicle for 48 hours in feeder free culture. Cell counting was carried out using CytoSmart cell counter, and validated by manual counting (A). Colony numbers per field were estimated under the microscope (B; 5 fields taken for each well, for 4 wells). The expression of *IRE1*, *ATF6* and *ASP* was examined by RT-qPCR following D6D inhibition for 24 hours (C). Apoptosis was assessed in TSCs following D6D inhibition for 24 hours using Annexin V/PI staining (D). The numbers in each quadrant represent the percentage of cells. Cell cycle was analyzed following inhibition of D6D by SC-26196 for 24 hours (E). n = 4; Data are presented as mean ± SEM. *, $P < 0.05$; **, $P < 0.01$; ***, $P < 0.001$.

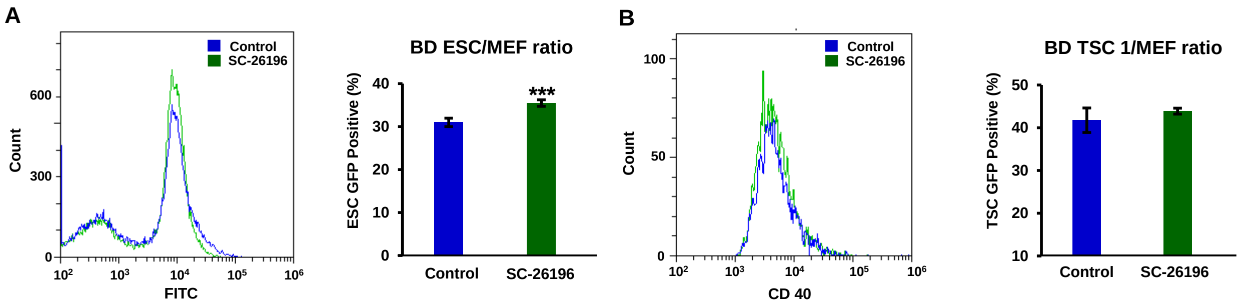


**Figure S3: Influence of D6D on SC proliferation rate in feeder cell-based cultures**. FACS analyses were carried out to assess the proliferation of ESCs and TSCs in co-culture with MEF feeder cells. The proliferation rate of ESCs and TSCs treated with SC-26196 (A-B; 0.2 µM) for 48 hours, was compared to their DMSO control as a ratio of SC to MEF number in culture. ESCs proliferation was assessed as a ratio of *Oct4*-GFP labeled cells to non-labeled cells (A); TSCs proliferation was assessed as a ratio of CD40-labeled cells to non-labeled cells (B). n = 4. Data are presented as mean ± SEM. *, $P < 0.05$; **, $P < 0.01$; ***, $P < 0.001$.

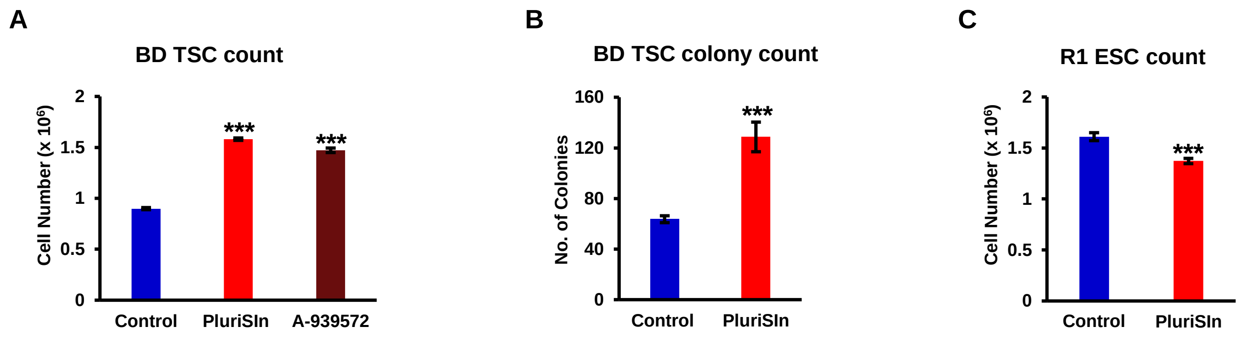


**Figure S4: The increase in viability of TSCs, whereas decrease of ESC viability by Scd1 inhibition is not cell line specific.** Following treatment with Scd1 specific inhibitor (PluriSIn 1; 20 µM), or DMSO vehicle, for 48 hours in feeder free culture, cell counting was carried out using CytoSmart cell counter, and further validated by manual counting of TSCs (Homo BL6 background; A) and ESCs (R1; C). Colony numbers per field were estimated under the microscope (B; 5 fields taken for each well, for 4 wells). n = 4. Data are presented as mean ± SEM. *, $P < 0.05$; **, $P < 0.01$; ***, $P < 0.001$.

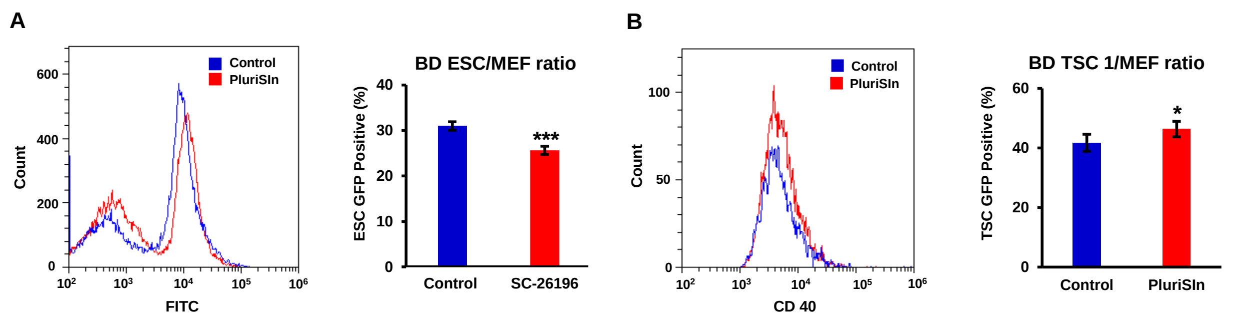


**Figure S5: The influence of Scd1 on SC proliferation rate in feeder cell-based culture**. FACS analysis was used to quantify the proliferation of ESCs and TSCs (ESCs and TSCs of the same background (129 / black 6) in co-culture with MEF feeder cells. The proliferation rate of ESCs (A) and TSCs (B) treated with PluriSIn 1 (20 µM) for 48 hours, was compared to their DMSO control as a ratio to MEFs in culture. ESCs proliferation was assessed as a ratio of *Oct4*-GFP labeled cells to non-labeled cells; TSCs proliferation was assessed as a ratio of CD40-labeled cells to non-labeled cells. n = 4. Data are presented as mean ± SEM. *, $P < 0.05$; **, $P < 0.01$; ***, $P < 0.001$.

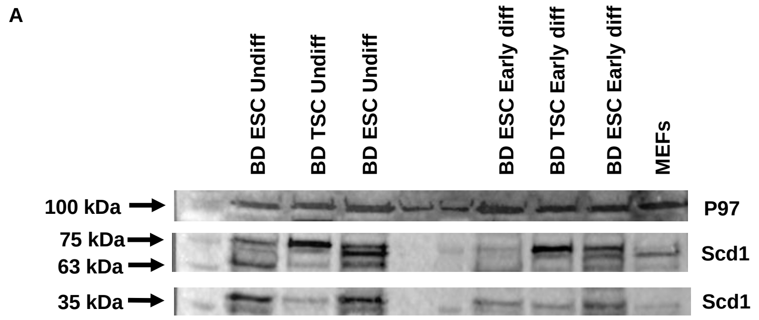


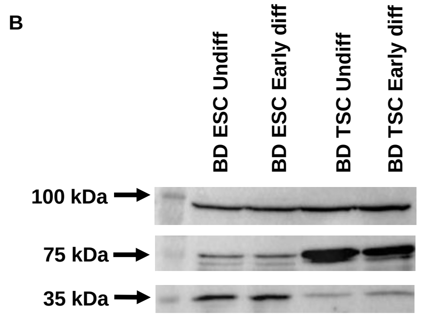


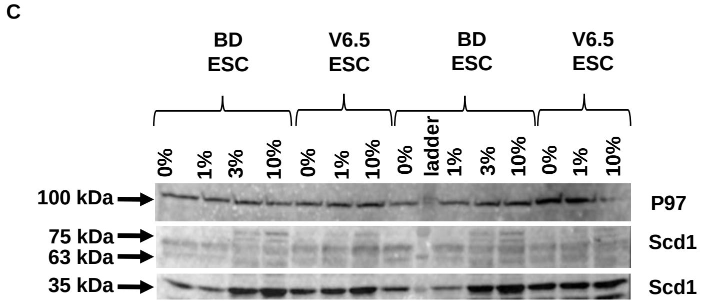


**Figure S6: Scd1 protein is expressed as a 37 kDa variant in ESCs, whereas largely expressed as a 70 kDa in TSCs.** BD ESCs and TSCs were incubated in ESC medium or Tx medium, respectively (A), or both in Tx medium (B), and the expression of Scd1 was assessed by Western blot analysis. BD ESCs and V6.5 ESCs were incubated in medium containing 0-10% fetal bovine serum for 48 hours. Scd1 protein expression was assessed by Western blotting (C).

**Original uncropped western blot images**

Uncropped western blot for the supplementary figure S6-A

Scd1

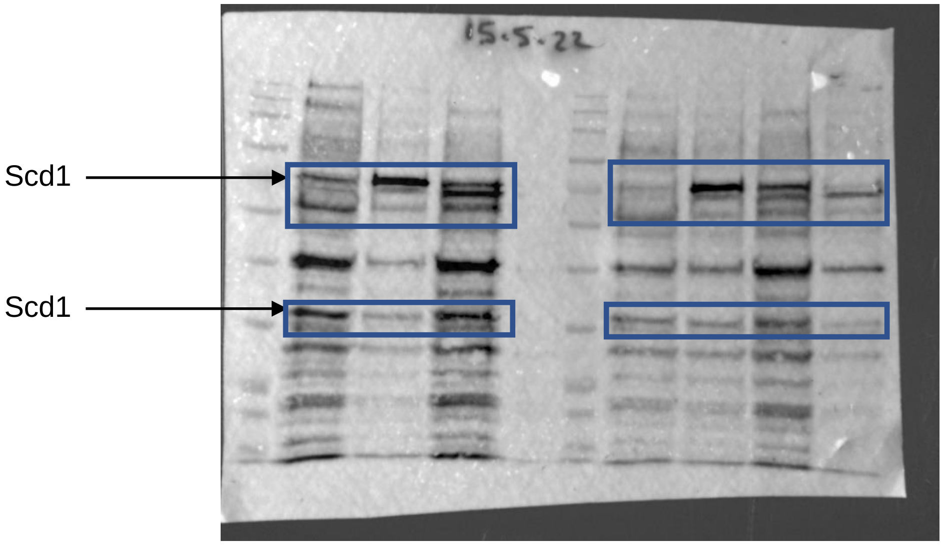


P97 (Loading control)

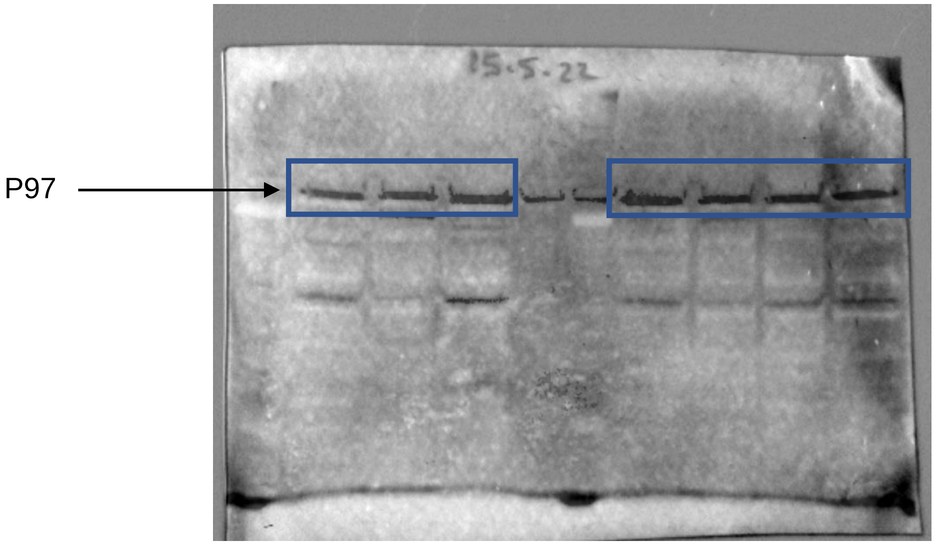

Uncropped western blot for the Supplementary Figure S6-B

Scd1

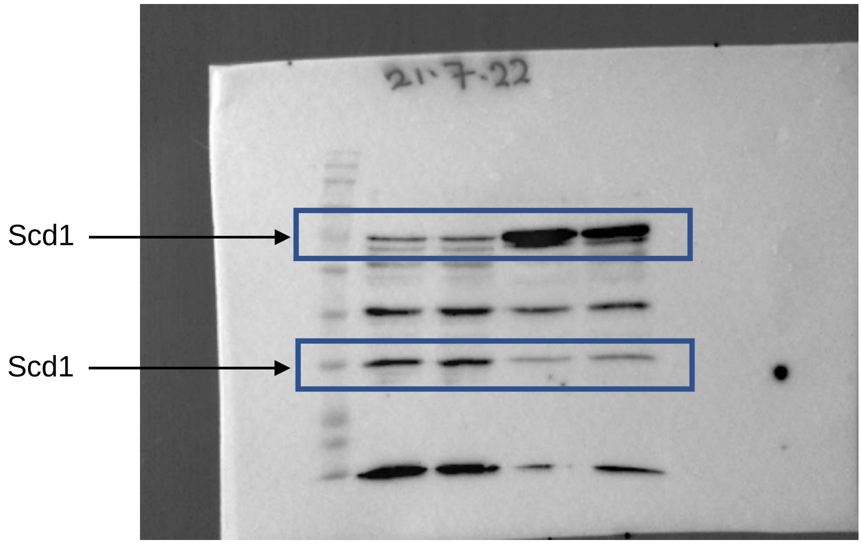


P97 (Loading control)

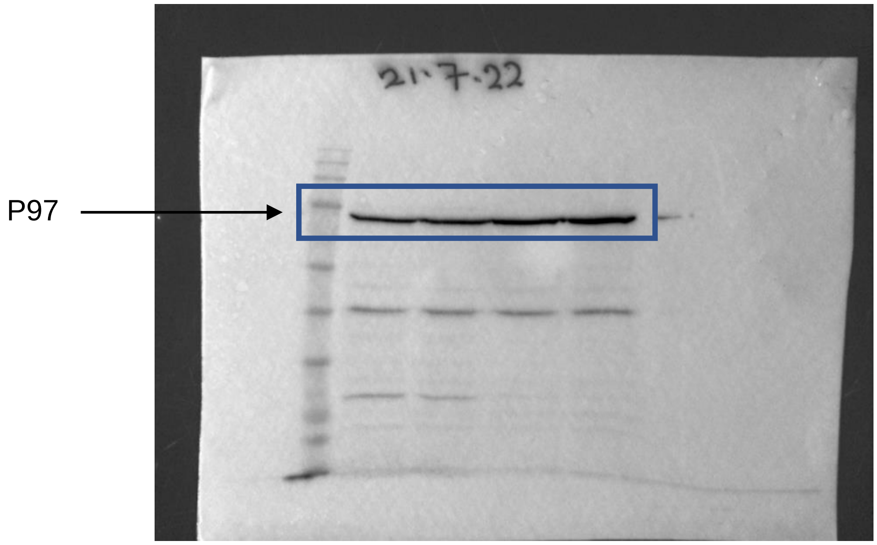

Uncropped western blot for the Supplementary Figure S6-C

Scd1

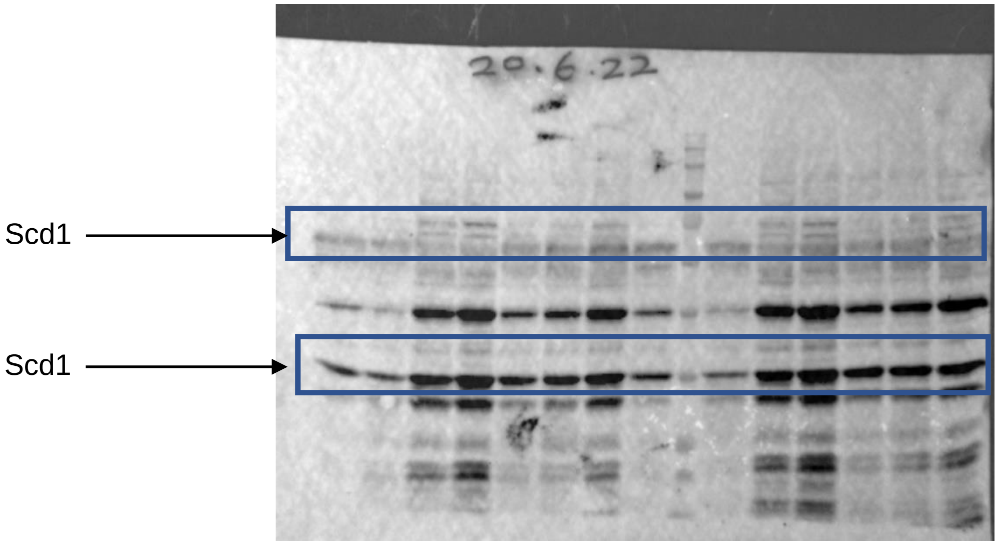


P97 (Loading control)

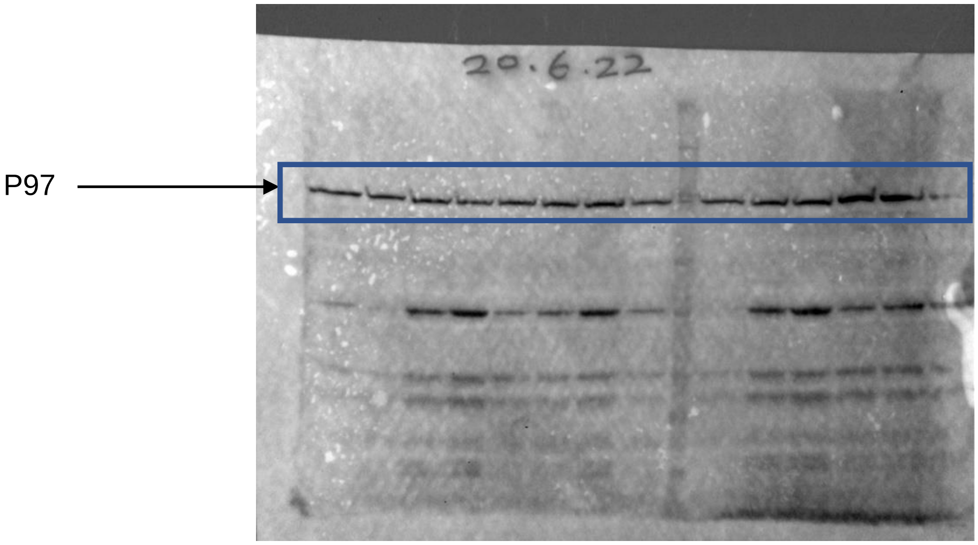

**Table S1: Discriminant lipid species identified in the early differentiated ESCs and TSCs**

| Lipid Species | Identification | Empirical Formula | m/z |
|---|---|---|---|
| 1 | PE(22:5/P-16:0) | $C_{43}H_{76}NO_7P$ | 748.52 (M-H) |
| 2 | PC(16:0/20:3) | $C_{43}H_{80}NO_8P$ | 828.57 (M+FA-H) |
| 3 | PE(P-16:0/18:4) | $C_{39}H_{70}NO_8P$ | 694.48 (M-H) |
| 4 | PI(20:4/18:0) | $C_{47}H_{83}O_{13}P$ | 885.54 (M-H) |
| 5 | PE(20:3/P-18:0) | $C_{45}H_{82}NO_7P$ | 838.59 (M+FA-H) |
| 6 | PI(20:3/18:0) | $C_{47}H_{85}O_{13}P$ | 887.56 (M-H) |
| 7 | PE(18:0/20:4) | $C_{43}H_{78}NO_8P$ | 766.53 (M-H) |
| 8 | PE(20:4/P-18:0) | $C_{43}H_{78}NO_7P$ | 750.52 (M-H) |
| 9 | PE(22:4/P-18:0) | $C_{43}H_{80}NO_7P$ | 776.56 (M-H) |
| 10 | PC(16:0/20:4) | $C_{44}H_{80}NO_8P$ | 826.55 (M+FA-H) |
| 11 | PE(22:6/18:0) | $C_{45}H_{78}NO_8P$ | 790.53 (M-H) |
| 12 | PE(16:0/22:6) | $C_{43}H_{74}NO_8P$ | 762.50 (M-H) |
| 13 | PE(20:5/18:1) | $C_{43}H_{74}NO_8P$ | 762.50 (M-H) |
| 14 | PE(22:6/P-16:0) | $C_{43}H_{74}NO_8P$ | 746.51 (M-H) |
| 15 | PG (20:4/18:3) | $C_{44}H_{73}O_{10}P$ | 837.49 (M+FA-H) |
| 16 | PE(22:6/16:1) | $C_{43}H_{72}NO_8P$ | 760.49 (M-H) |
| 17 | PG(18:1/22:5) | $C_{46}H_{79}O_{10}P$ | 821.54 (M-H) |
| 18 | PE(P-18:0/22:6) | $C_{45}H_{78}NO_7P$ | 774.54 (M-H) |
| 19 | PE(16:1/20:3) | $C_{41}H_{74}NO_8P$ | 738.50 (M-H) |
| 20 | PI (18:1/18:1) | $C_{45}H_{83}O_{13}P$ | 861.54 (M-H) |
| 21 | PG (16:1/18:1) | $C_{40}H_{75}O_{10}P$ | 745.50 (M-H) |
| 22 | PC(14:0/18:1) | $C_{40}H_{78}NO_8P$ | 1508.09 (2M+FA-H) |
| 23 | PE(18:1/18:1) | $C_{23}H_{46}NO_7P$ | 742.53 (M-H) |
| 24 | SM(18:1/16:0) | $C_{39}H_{79}N_2O_6P$ | 747.56 (M+FA-H) |
| 25 | PG(18:0/18:1) | $C_{42}H_{81}O_{10}P$ | 775.54 (M-H) |
| 26 | PE(P-18:1/18:1) | $C_{41}H_{78}NO_7P$ | 726.54 (M-H) |
| 27 | PE(18:1/16:0) | $C_{39}H_{76}NO_8P$ | 716.52 (M-H) |
| 28 | PE(18:1/16:1) | $C_{39}H_{74}NO_8P$ | 715.51 (M-H) |
| 29 | PE(18:1/20:1) | $C_{43}H_{82}NO_8P$ | 770.56 (M-H) |
| 30 | PC(14:0/16:1) | $C_{38}H_{74}NO_8P$ | 748.54 (M+FA-H) |
| 31 | PG(18:1/18:1) | $C_{42}H_{78}O_{10}P$ | 773.53 (M-H) |
| 32 | Phospholipid (34:2)* | $C_{39}H_{74}NO_8P$ | 715.50 (M-H) |

| | | | |
|---|---|---|---|
| 33 | Phospholipid (34:1)* | $C_{41}H_{80}NO_8P$ | 744.54 (M-H) |
| 34 | Phospholipid (34:1)* | $C_{39}H_{76}NO_8P$ | 716.52 (M-H) |
| 35 | Phospholipid (36:3)* | $C_{42}H_{77}O_{10}P$ | 771.51 (M-H) |
| 36 | Phospholipid (40:6)* | $C_{46}H_{80}NO_8P$ | 850.55 (M+FA-H) |
| 37 | Phospholipid (34:3)* | $C_{42}H_{78}NO_8P$ | 800.54 (M+FA-H) |
| 38 | Phospholipid (38:4)* | $C_{44}H_{79}O_{10}P$ | 797.53 (M-H) |
| 39 | Phospholipid (36:1)* | $C_{44}H_{86}NO_8P$ | 832.60 (M+FA-H) |

PE, Phosphatidylethanolamine; PC, Phosphatidylcholine; PI, Phosphatidylinositol; PG, Phosphatidylglycerol; SM, Sphingomyelin. The lipids were identified by $MS^E$. * Identified as phospholipids based on diagnostic ions (no full assignment).

**Table S2: Temperature gradient for Gas chromatography**

| Rate (°C/min) | Value (°C) | Hold Time (Min) | Run Time (Min) |
|---|---|---|---|
| | 50 | 5 | 5 |
| 10 | 170 | 0 | 17 |
| 3.5 | 250 | 3 | 42 |

**Table S3: List of primers used for RT-qPCR**

| Mouse Genes | Forward Primer (Sequence: 5' to 3') | Reverse Primer (Sequence: 5' to 3') |
|---|---|---|
| ***Ubc*** | CAGCCGTATATCTTCCCAGACT | CTCAGAGGGATGCCAGTAATCTA |
| ***Hprt*** | ATTCACCCCCACTGAGACTG | AGGGCATATCCAACAACAAACTT |
| ***Scd1*** | TTCTTGCGATACACTCTGGTGC | CGGGATTGAATGTTCTTGTCGT |
| ***D6D1*** | TCATCGGACACTATTCGGGAG | GGGCCAGCTCACCAATCAG |
| ***D6D2*** | GACCGTGGCAAAAGCTCTCA | CTGTGACGAGGGTAGGAATCC |
| ***D5D*** | ACCCAGCTTTGAACCCACC | GAGGCCCATTCGCTCTACTG |
| ***Nanog*** | AAACCAGTGGTTGAAGACTAGCAA | GGTGCTGAGCCCTTCTGAATC |
| ***Oct4*** | GCTTGGGCTAGAGAAGGATGTG | TGGCGCCGGTTACAGAAC |
| ***Sox2*** | GCGGCGGAAAACCAAGA | CCGGGAAGCGTGTACTTATCC |
| ***Cdx2*** | CACTTTAGTCGATACATCACCATC | GATTTTCCTCTCCTTGGCTCT |
| ***Elf5*** | GGACCGATCTGTTCAGCAAT | GGGTGCACTGATGTCCAGTA |
| ***Tfap2c*** | CAGGTCACTCTCCTCACGTC | CAGCTTCGCAGACATAGGC |
| ***Eomes*** | CGGCAAAGCGGACAATAACA | GGAGCCAGTGTTAGGAGATTC |
| ***Asp*** | CTAGCCATCACTTATGGTTTGGT | TGCCCAGATTACTAAACTGCCA |
| ***Atf3*** | CCCCTGGAGATGTCAGTCAC | CGGTGCAGGTTGAGCATGT |
| ***Atf6*** | CAGTTGCTCCATCTCCTCTCC | TGGGACACTGGCATTGGTTTG |
| ***Ire1α*** | GAGCAAGCTAACGCCTACTCTGT | CACCATTGAGGGAGAGGCATA |
